# A systematic and quantitative comparison of lattice and Gaussian light-sheets

**DOI:** 10.1101/2020.06.12.147181

**Authors:** Bo-Jui Chang, Kevin M. Dean, Reto Fiolka

## Abstract

The axial resolving power of a light-sheet microscope is determined by the thickness of the illumination beam and the numerical aperture of its detection optics. Bessel-based optical lattices have generated significant interest owing to their potentially narrow beam waist and propagation-invariant characteristics. Yet, despite their significant use in Lattice Light-Sheet Microscopy, and recent incorporation into commercialized systems, there are very few quantitative reports on their physical properties and how they compare to standard Gaussian illumination beams. Here, we systematically measure the beam properties in transmission of dithered square lattices, which is the most commonly used variant of Lattice Light-Sheet Microscopy, and Gaussian-based light-sheets. After a systematic analysis, we find that square lattices are very similar to Gaussian-based light-sheets in terms of thickness, confocal parameter and propagation length.

## 1. Introduction

Light-sheet fluorescence microscopy (LSFM) is a rapidly growing volumetric imaging technique that offers life scientists a unique combination of speed, sensitivity, optical sectioning, and low phototoxicity and photobleaching [1]. In LSFM, a biological specimen is illuminated with a focused sheet of light, and fluorescence originating from this sheet of light is captured in a widefield format using modern scientific cameras. Thus, unlike traditional epi-fluorescence imaging methods that nonproductively illuminate regions above and below the detection objective depth-of-focus, LSFM restricts the potentially damaging illumination to only the in-focus region of the specimen. This not only decreases phototoxicity and photobleaching, but delivers higher contrast imaging owing to its inherent optical sectioning capability [2].

Several factors critically influence the performance of LSFM [3]. Like conventional epifluorescence widefield microscopes, the lateral resolution of LSFM scales linearly with the fluorescence emission wavelength (*λ*_*EM*_) and inversely with the numerical aperture (*NA_DET_*) of the detection objective, *λ*_*EM*_/2*NA_DET_*. However, the axial resolution of LSFM depends upon the wavelengths of the illumination sheet (*λ*_*EX*_) and fluorescence emission (*λ*_*EM*_), and the numerical aperture of the excitation light-sheet (*NA_EX_*) and detection objective (*NA_DET_*). In the Fourier domain, this can be estimated from the highest axial frequencies that contribute to image formation, where θ_*DET*_ is the solid angle over which the detection objective collects light and *n* is the refractive index of the medium. As a result, the axial as.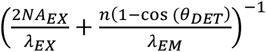 [3]. Of note, this is an optimistic estimate that assumes no attenuation of higher axial frequencies (e.g., optical transfer function roll-off), but it nevertheless serves as an upper bound for the axial resolution of LSFM. Consequently, for thick light-sheets, the numerical aperture of the detection objective largely dictates the axial resolution of the imaging system. In contrast, if the light-sheet can be made thinner than the detection objective depth of focus, the axial resolution approaches that of the illumination beam waist.

Critically, for Gaussian optics, the field of view of LSFM is only as large as the region over which the illumination beam approximates a sheet of light, which can be estimated as ~2 Rayleigh lengths 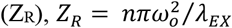, and is also referred to as the confocal parameter. However, the Rayleigh length, and hence the field of view, decreases nonlinearly as the beam waist, *w*_*o*_, becomes more tightly focused. Hence, one must make a critical trade-off between the field of view and the thickness of the illumination beam.

In an effort to overcome the tradeoff between the size of the field of view and light-sheet thickness, several labs adopted propagation invariant illumination beams (e.g., Bessel and Airy beams), which in theory, maintain a tight 2D focus throughout an infinite propagation length [4–7]. In practice, such beams are finite in length and accompanied with sidelobes structures that grow in magnitude with the propagation length of the beam [8]. For many cases, these sidelobe structures contain more excitation energy than the tightly focused central lobe, which results in heightened image blur and diminished optical sectioning. As such, in an effort to partially reduce blur and restore image contrast, iterative deconvolution routines become increasingly necessary. Thus, while propagation invariant beams overcame the divergence of Gaussian optics, they do so by compromising optical sectioning (which is a key feature of LSFMs), potentially increasing more photo-bleaching and introducing significant computational overhead.

More recently, carefully crafted coherent superpositions of Bessel beams have been reported to provide a compromise between light-sheet thickness and sidelobe energy density [9]. Two major variants of the technique, which is referred to as Lattice Light-Sheet Microscopy (LLSM), exist. 1) The hexagonal lattice, which has a narrow propagation invariant beam waist surrounded by a series of sidelobes that grow in strength with increasing propagation length, and 2) the square lattice, which consists of a thicker central beam lobe that grows in size with increasing propagation length but has small side lobes. Although the former provides superior axial resolution, the latter is the most widely adopted, as it achieves a more favorable balance between optical sectioning and axial resolution. Indeed, the square lattice has served as the bedrock for a number of stunning publications that reveal the beauty and complexity of biological systems [10–13].

While LLSM has received significant interest and has been hailed for its unique combination of gentle illumination, high-resolution, and high-speed imaging, many key optical parameters have not been independently investigated. These include the light-sheet thickness and the confocal parameter, which defines the distance over which a light-sheet remains tightly focused and thus pragmatically useful. Indeed, a recent study based on numerical simulations challenged the notion that the optical lattices adopted by LLSM, in particular the square lattice, are superior to Gaussian light-sheets [14]. Nonetheless, owing to the complexities in modeling high NA optical lattices, we saw a strong need to verify these predictions experimentally.

In this paper, we set out to experimentally compare the optical characteristics of dithered square lattices, which are most commonly used in LLSM (for example, see Supporting Table S1 in [15]), and Gaussian beams. Importantly, we find that Gaussian beams and square optical lattices share surprisingly similar optical properties, including beam waist thickness and confocal parameter. Thus, we stipulate that LSFMs equipped with Gaussian beams will perform identically to a LLSM operating with a dithered square lattice. Furthermore, we find that at the numerical apertures used here, which are needed for high axial resolution, the usual approximation for Gaussian light-sheets is inaccurate. And lastly, we explore different LLSM tuning parameters that have not been discussed in detail in the literature and document how they affect light-sheet performance.

## 2. Materials and Methods

### 2.1 Experimental setup for Light-Sheet generation

A detailed schematic of the experimental setup is shown in Appendix A. The system allows the generation of optical lattices and Gaussian light-sheets and possesses a detection path with which these light-sheets can be imaged in transmission. In general, all lenses were placed in a 4f arrangement, which was confirmed with a shear plate interferometer, and the lateral and rotational alignment of each element was confirmed by measuring residual back-reflections from each optical surface. For all experiments described here, a 488-nm continuous wave laser (Sapphire 488-300 LP, Coherent) was used as the light source, which was spatially filtered by focusing it through a 50-μm pinhole (P50D, Thorlabs) with a 50-mm focal length achromatic lens (AC254-050-A, Thorlabs) prior to being diverted down one of two optical paths with a flip mirror.

The first path is a modified version of a LLSM [9], which has been described in detail elsewhere [15]. Here, the laser is recollimated with a 75-mm achromatic doublet (AC254-075-A, Thorlabs) prior to asymmetric expansion into a line profile with a pair of achromatic cylindrical lenses (68-160, Edmund Optics, and ACY254-200-A, Thorlabs). This line-shaped beam uniformly illuminates a narrow strip on a spatial light modulator (SLM, SXGA-3DM, Forth Dimension Displays) where binary phase holograms are displayed to generate the desired optical lattice. A polarized beam splitter (10FC16PB.3, Newport) and a half-wave plate (AHWP10M-600, Thorlabs) are placed in front of the SLM, and together with the SLM, form a reconfigurable lattice light-sheet generator. The polarization of the input laser light was adjusted to maximize the light-throughput through this unit. The diffracted light from the SLM is focused by an achromatic lens (AC508-400-A, Thorlabs) onto a custom-designed binary transmission mask (Photo Sciences, Inc.), which consists of multiple annuli of various sizes that serve to block the unwanted zeroth and higher-order (e.g., N > 1) diffracted beams, and to specify the inner and outer numerical apertures of the lattice light-sheet. After passing through the mask, the desired diffraction orders are de-magnified through two achromatic lenses (AC254-150-A and AC254-100-A, Thorlabs) and projected onto a mirror galvanometer (6215H, Cambridge Technology) for rapid lateral dithering of the illumination light-sheet. Thereafter, the light reflects off of a folding mirror, and is relayed to the back pupil of the 40X NA 0.8 illumination objective (CFI Apo 40XW NIR, Nikon Instruments) with an achromatic lens (AC-254-100-A, Thorlabs), a second flip mirror, and a tube lens (ITL-200, Thorlabs).

The second optical path is recollimated with an 80-mm achromatic doublet (AC254-080-A, Thorlabs), and is used to create conventional Gaussian-based illumination light-sheets. Here, the laser light is asymmetrically magnified into a line profile by a pair of achromatic cylindrical lenses (ACY254-050-A and ACY254-200-A, Thorlabs), and focused with a cylindrical lens (ACY254-050-A, Thorlabs) onto a mirror galvanometer (6215H, Cambridge Technology) that is conjugate to the focal plane of the illumination objective for rapid light-sheet pivoting [16]. Thereafter, the light is reflected with a folding mirror, and relayed to the back focal plane of the illumination objective with an achromatic lens (AC254-075-A, Thorlabs) and two tube lenses (ITL-200, Thorlabs). An adjustable slit, which is placed conjugate to the back focal plane of the illumination objective, is used to control the numerical aperture of the Gaussian illumination light-sheet. Besides the slit, we also used annular masks and placed the line beam at specific position to generate Gaussian light-sheets (see also Appendix B). We used these datasets to supplement our light-sheet measurements with the variable slit. We note that in either way (slit or annular mask), only the central flat-top portion of the Gaussian input beam remains: I.e. the intensity profile in the back focal plane is relatively uniform, instead of possessing a Gaussian profile. But we decided to use the term Gaussian light-sheet throughout the manuscript, as this is a very common terminology in the light-sheet field.

Both the LLSM and conventional Gaussian-based light-sheets were evaluated with a camera conjugate to the back pupil of the illumination objective. Here, a third flip mirror directs the laser light towards the camera (DCC1545M, Thorlabs), and two achromatic doublets (AC254-100-A and AC254-050-1, Thorlabs), and a neutral density filter (ND10A, or ND20A, Thorlabs), served to relay and attenuate the laser power, respectively. For LLSM, the correct placement of the diffraction orders in relation to the annular mask was monitored and the inner and outer numerical apertures of the illumination beam were measured. For conventional Gaussian-based LSFM, the width of the line profile in the back focal plane was measured to determine the numerical aperture of the illumination.

### 2.2 Light-Sheet Measurements

To measure the characteristics of the light-sheets in the front focal plane of the illumination objective, the light-sheets were imaged in a direct transmission mode using another 40X NA 0.8 objective (CFI Apo 40XW NIR, Nikon Instruments). Here, a 400-mm focal length achromatic lens (AC508-400-A) served as the tube lens, and the laser light was detected with a scientific CMOS camera (Orca Flash 4.0 V2, Hamamatsu Corp.) after attenuation with a neutral density filter (ND40A, Thorlabs). The combination of a 400-mm tube lens and the 40X detection objective offers a final magnification of 80X, which results in a pixel size of 81.25 nm. To acquire three-dimensional data sets of the light-sheets, the detection objective was scanned along the direction of laser propagation with a piezoelectric stage (P621.1CD, Physik Instrumente). The data acquired encompassed the main in-focus region of the light-sheet, as well as the more diffuse out-of-focus regions before and after it (See Figure 1).

**Fig. 1.**
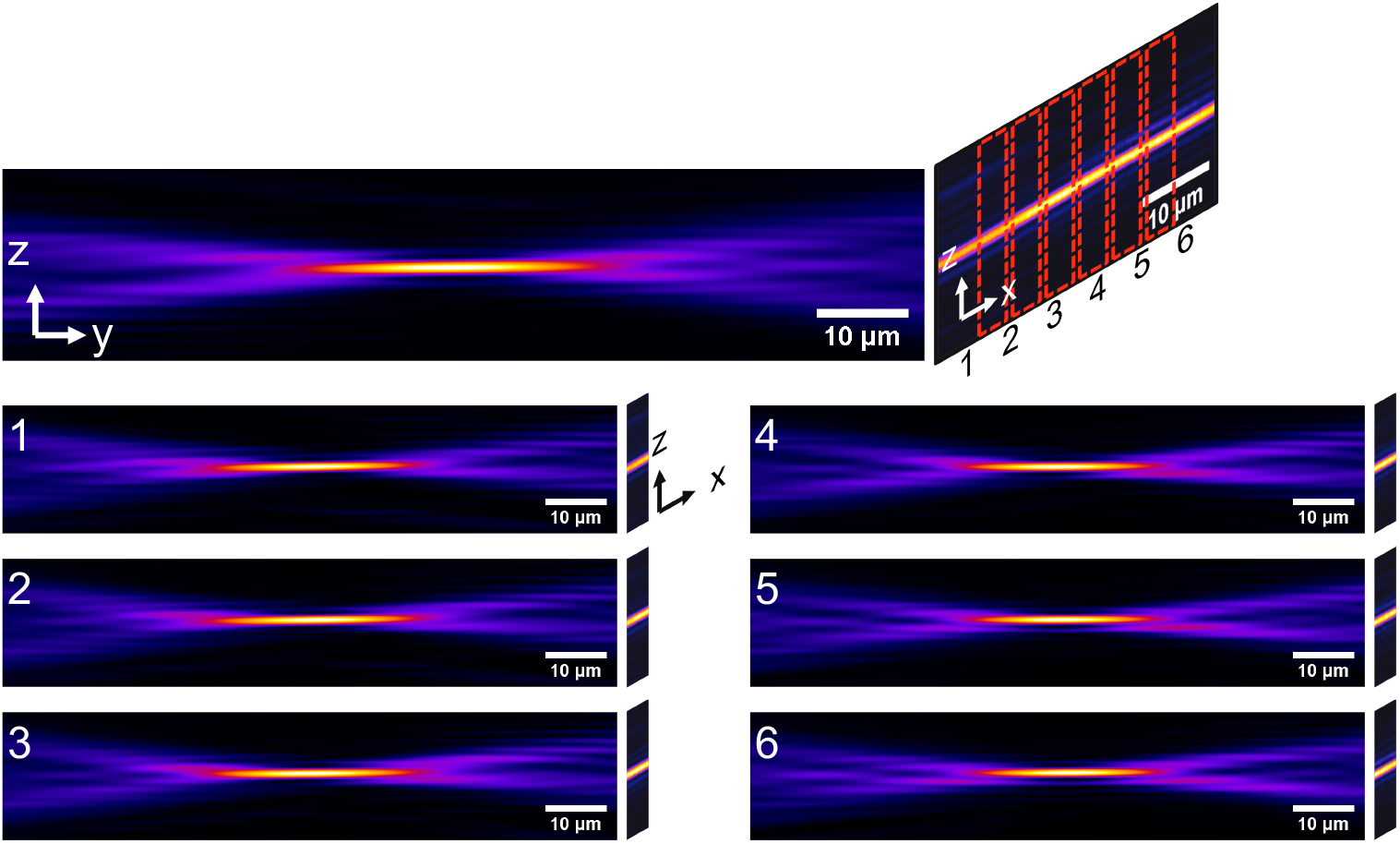
Characterization of light-sheets. The top panel represents a 3D stack of one Gaussian light-sheet. We subdivided the light-sheet in the transverse plane (x) into 6 individual blocks, numbered 1-6, which are shown below. In each of these blocks, the evolution of the beam waist along the propagation direction (y) was analyzed individually. The Gaussian light-sheet presented here correspond to a numerical aperture of approximately 0.2.

Importantly, it should be noted that all optical imaging systems generate a slightly blurred image of the real object (here, the light-sheets), which also holds for our transmission imaging setup. However, because the light-sheets that we have measured have a lower numerical aperture (0.19 – 0.64) than our detection objective (0.80) and hence were significantly thicker than its diffraction limit, we expect this effect to be negligible. Furthermore, both the Gaussian and LLSM light-sheets were measured identically, and thus they are subject to the same amount of optical blurring. Lastly, by measuring the light-sheets in transmission, we argue that this avoids potential challenges that could arise from measuring the intensity distribution of the light-sheet with sub-diffraction fluorescent nanospheres (e.g., bead aggregation, saturation, bleaching etc.).

### 2.3. Light-Sheet Analysis

All light-sheet measurements were performed in a fully automatic fashion using a publicly available software written in MATLAB (https://github.com/AdvancedImagingUTSW). Since the light-sheet stays constant over hundreds of microns in the transverse direction (labeled X in Figure 1, we subdivided the acquired data into 6 sub-volumes along the X-direction and analyzed each block individually. This allowed us to confirm that the light-sheet was uniform in the X-direction and allowed for averaging of the measured parameters.

For each light-sheet, three parameters are measured: 1) The Full-Width Half-Maximum (FWHM) of the light-sheet thickness in the Z-direction at the waist of the beam. 2) The FWHM of the light-sheet propagation length in the Y-direction. 3) The confocal parameter, for which we measure the distance between the two points along the Y-direction where the light-sheet thickness increases by a factor of 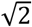 from its smallest value.

To measure the light-sheet thickness, we first evaluated each XZ plane along the Y-axis to find the focus of the light-sheet, i.e. the location of the highest beam intensity (Figure 2A). Next, the image dataset was rotated (usually less than 1°) such that the light-sheet was perfectly aligned perpendicular to the Z-axis (Figure 2B). Figure 2C shows a line profile of the intensity along the Z-axis along the center of the light-sheet. We measured the light-sheet thickness from the standard deviation (σ) of a Gaussian function fit 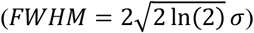. Importantly, a Gaussian function might be a poor approximation for light-sheets with strong sidelobes (e.g., Bessel and Hexagonal lattices) [14]. However, we found that at the beam waist, the profile was well approximated by a Gaussian function for both the Gaussian light-sheets as well as the square lattice light-sheets (Figure 2C).

**Fig. 2.**
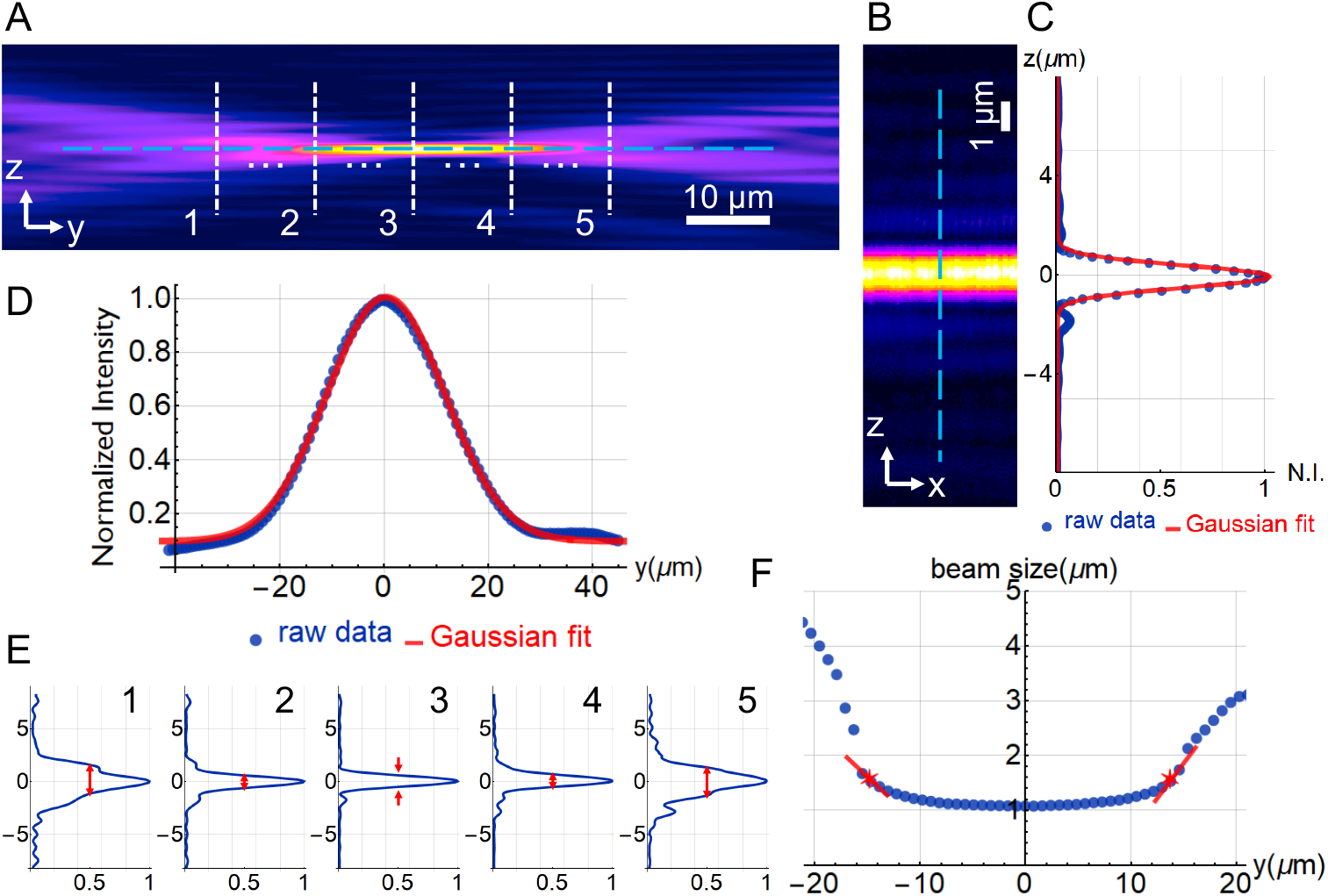
Details of the light-sheet quantification. (A) An YZ image of the light-sheet averaged along the X-axis. (B) XZ image at the waist of the light sheet. (C) The Z-intensity profile along the blue-dashed line shown in (B), and a Gaussian fit. The FWHM of the fitted curve is used to represent the thickness of the light-sheet. (D) The intensity profile of the blue dashed line in (A) and a Gaussian fit. The FWHM of the fitted curve is used to represent the propagation length of the light-sheet. (E) Intensity profiles of the five dashed white lines in (A). (F) The beam size along the propagation direction (Y) of the light-sheet. The red lines go through the points that are close to 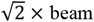 waist, and the star points show exactly where the beam size increased to 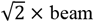 waist by interpolation along the red lines. The distance between the star points represents the confocal parameter. The data set shown here is a Gaussian light-sheet generated with an NA of approximately 0.2.

To measure the propagation length, we first average the plane-by-plane image (YZ slice) along the X-axis for each block, and then we rotate the image slightly (usually also less than 1°) to make sure the propagation of the light-sheet is parallel to the Y-axis. Figure 2D shows a line profile of the intensity along the Y-axis along the center of the light-sheet. We measured the beam length from the standard deviation (σ) from the fit of a Gaussian function in the same way as we do for the light-sheet thickness.

To measure the confocal parameter of the light-sheet, we measured the width of the light sheet in the Z-direction at multiple positions along the Y-axis, *i.e.* the intensity profiles of the white dashed lines in Figure 2A. Although a Gaussian curve fit provided satisfactory results near the beam waist, it failed in regions away from the light-sheet focus as the intensity distribution becomes increasingly non-Gaussian. Thus, we numerically estimated the FWHM of the light-sheet thickness by interpolating where the intensity drops to 50% of its peak value (Figure 2E). As can be seen in Figure 2F, the light-sheet thickness remained relatively constant over a length ~20 microns but grows rapidly outside of this range. From this curve, we then numerically identified the confocal parameter, which is defined as two times the distance over which it takes the light-sheet thickness to increase by a factor of 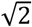 (red stars in Figure 2F). We wanted to emphasize that for a purely Gaussian beam, the confocal parameter would be twice the Rayleigh length, which can be computed analytically from the beam waist thickness. However, we have observed that our light-sheets do not behave exactly like Gaussian light-sheets, i.e. their beam waist stays constant in the main lobe of the light-sheet, and then increases rather rapidly outside, yielding poor results if fitted with a quadratic function. As such, we decided to report the confocal parameter as defined by the 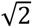 thickness increase.

### 2.4 Light-Sheet Simulations

For numerical simulations of Gaussian and square lattices light-sheets, we have used the publicly available code from *Remacha et al* (see https://github.com/remachae/beamsimulator) [14]. For all simulations, a refractive index of *n* = 1.33 and an excitation wavelength of *λ*_*EX*_ = 0.488 *μm* was assumed. We decided to use their definition of main-lobe thickness (*W*_*ML*_) and Length, which are output parameters of the simulation. As previously noted, the square lattice contains small sidelobes, but they are small enough that *W*_*ML*_ can effectively capture the thickness of it. *W*_*ML*_ is defined as the beam size when the intensity drops by a factor of 1/*e*. Thus, the factor *FWHM* ≅ 0.8326 × *W*_*ML*_ is applied to the simulation results. We also include the theoretical values for Gaussian light-sheets using an analytical formula for Gaussian beams [17]. The beam waist, *w*_*o*_, used in Gaussian Optics is the radius of the beam when the intensity drops to 1/*e*^2^, and relates to the FWHM in the following way: *w*_*o*_ ≅ 0.85 × *FWHM*. By applying these relationships, the thickness (given as FWHM) and the confocal parameter (2*Z*_*R*_) are obtained.

## 3. Results

### 3.1. Systematic Evaluation of Light-Sheet Properties

To evaluate the optical properties of the square lattice light-sheet in LLSM, we systematically varied the inner (na) and outer (NA) numerical apertures of the illumination annulus and measured the thickness and propagation length of the light-sheet in transmission. In each case, a matching hologram was applied to the SLM, and the correct diffraction pattern was confirmed with the camera that is conjugate to the back focal plane of the illumination objective. We further evaluated parameters linked to the hologram generation, which among other things influenced the relative positioning of the diffraction orders relative to the annular mask. Lattice light-sheets were rapidly dithered in the X-direction with a mirror galvanometer to eliminate the lateral interference patterns. To facilitate a side-by-side comparison, we report measurements on nearly all of the square lattice light-sheets from the original Lattice light sheet microscopy manuscript [9]. We then acquired an exhaustive series of Gaussian light-sheets where the excitation NA was varied in small steps. From this data set, Gaussian light-sheets were selected that closely matched their lattice light-sheet equivalents. A list of the NAs for the measured Gaussian and square lattice light-sheets is provided in the Appendix C.

Qualitative differences are readily visualized in the cross-sectional views for the dithered square lattice and Gaussian light-sheets (Figure 3). For example, the dithered square lattice, while having a similar shaped central lobe compared to the Gaussian sheet, exhibits some structural differences in the transition zones before and after the light-sheet focus (white arrows). In addition, the dithered square lattice features two distinct sidelobes (white arrows in XZ-view), whereas the Gaussian sheet is almost free of sidelobes. Further, in Figure 3A-B, one can see two faint beams coming from much steeper angles in the cross-sectional view for the square lattice, which are not present in the Gaussian sheet. These beams originate from the central diffraction order of the hologram used for lattice generation. Interestingly, we experimentally blocked these orders in Appendix D and found that they negligibly influence the properties of the light-sheet.

**Fig. 3.**
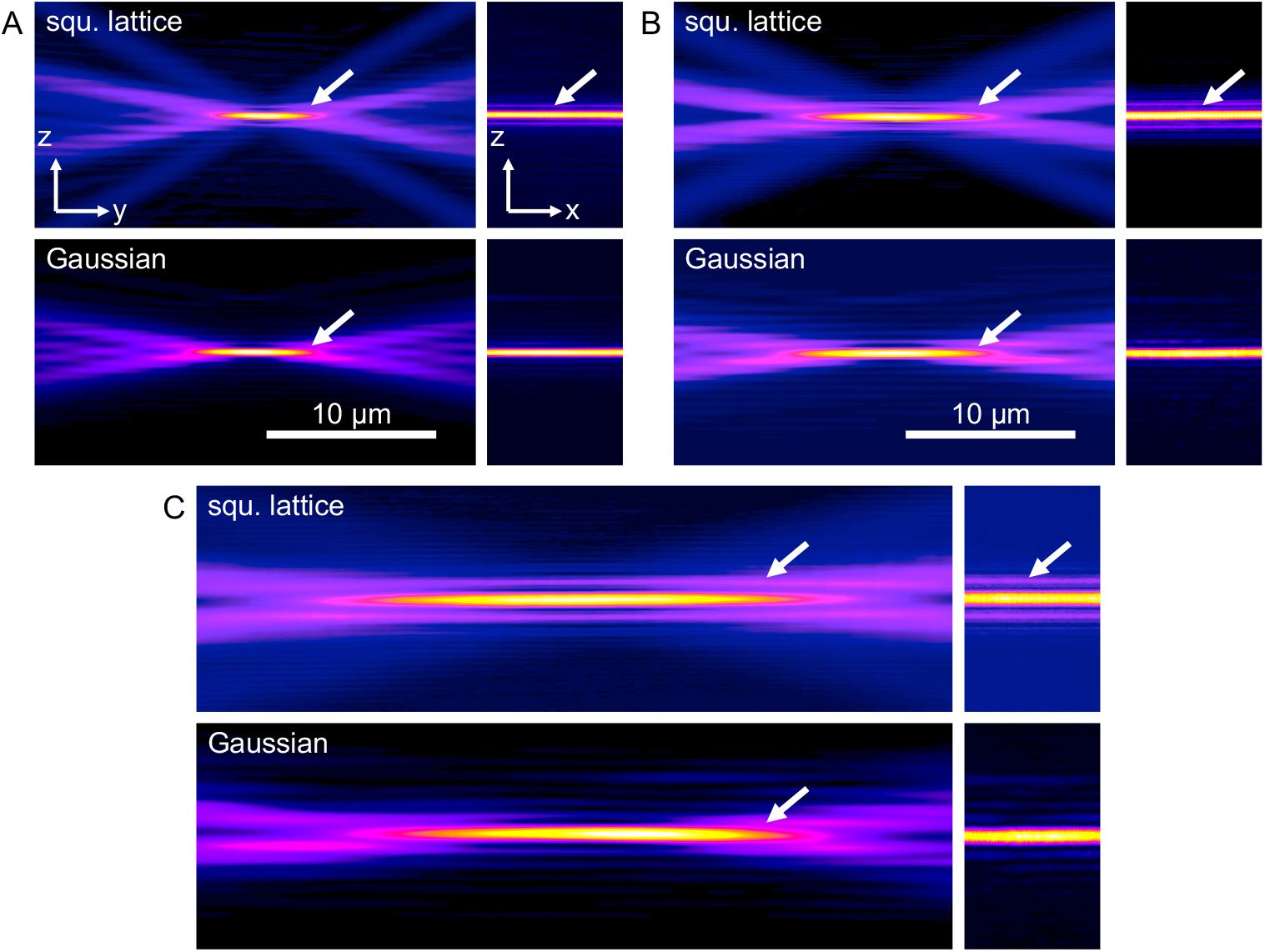
Qualitative comparison of dithered square lattice and Gaussian light-sheet. Square lattice light-sheets were generated with (A) NA 0.70 and na 0.56, (B) NA 0.55 and na 0.44, and (C) NA 0.35, na 0.28 annulus. Y-Z and X-Z cross-sectional views are shown. The X-Z view is located at the center (thinnest waist) of each light-sheet. The Gaussian light-sheet (NA= 0.31, 0.24, and 0.14 in (A), (B), (C), respectively) is chosen to have similar properties to each corresponding square lattice light-sheet. Arrows point at differences at the edges of the useful light-sheet and at sidelobes in the lattice light-sheet. NA: outer numerical NA of annulus used for squared lattice; na: inner numerical aperture for annulus used for the squared lattice. For Gaussian beams, NA refers to the effective illumination NA used for this beam.

In an effort to facilitate a more detailed assessment of light-sheet properties, Figure 4 provides detailed profiles for three square lattices and three Gaussian light-sheets with a matching confocal parameter. Strikingly, Figure 4A shows that both light sheets maintain a nearly constant thickness throughout their optical focus (e.g., within the range dictated by their confocal parameters). This is a property that was expected for the lattice light-sheet, as it derives from Bessel beams, which in turn have a constant intensity profile over a finite propagation length. It is however a surprise for the Gaussian light-sheets, which were expected to show a quadratic increase in beam waist thickness over their length. Importantly, the cross-sectional profiles (Figure 4B) show a close correspondence in their main lobe thickness, which influences the axial resolution of LSFM. The square lattice is accompanied with side lobes that grow in intensity with increasing confocal parameter, more dominantly so than for the Gaussian light-sheet. Thus, for a given light-sheet confocal parameter, Gaussian beams appear to provide similar axial resolution, albeit with reduced image blur and greater optical sectioning capability.

**Fig. 4.**
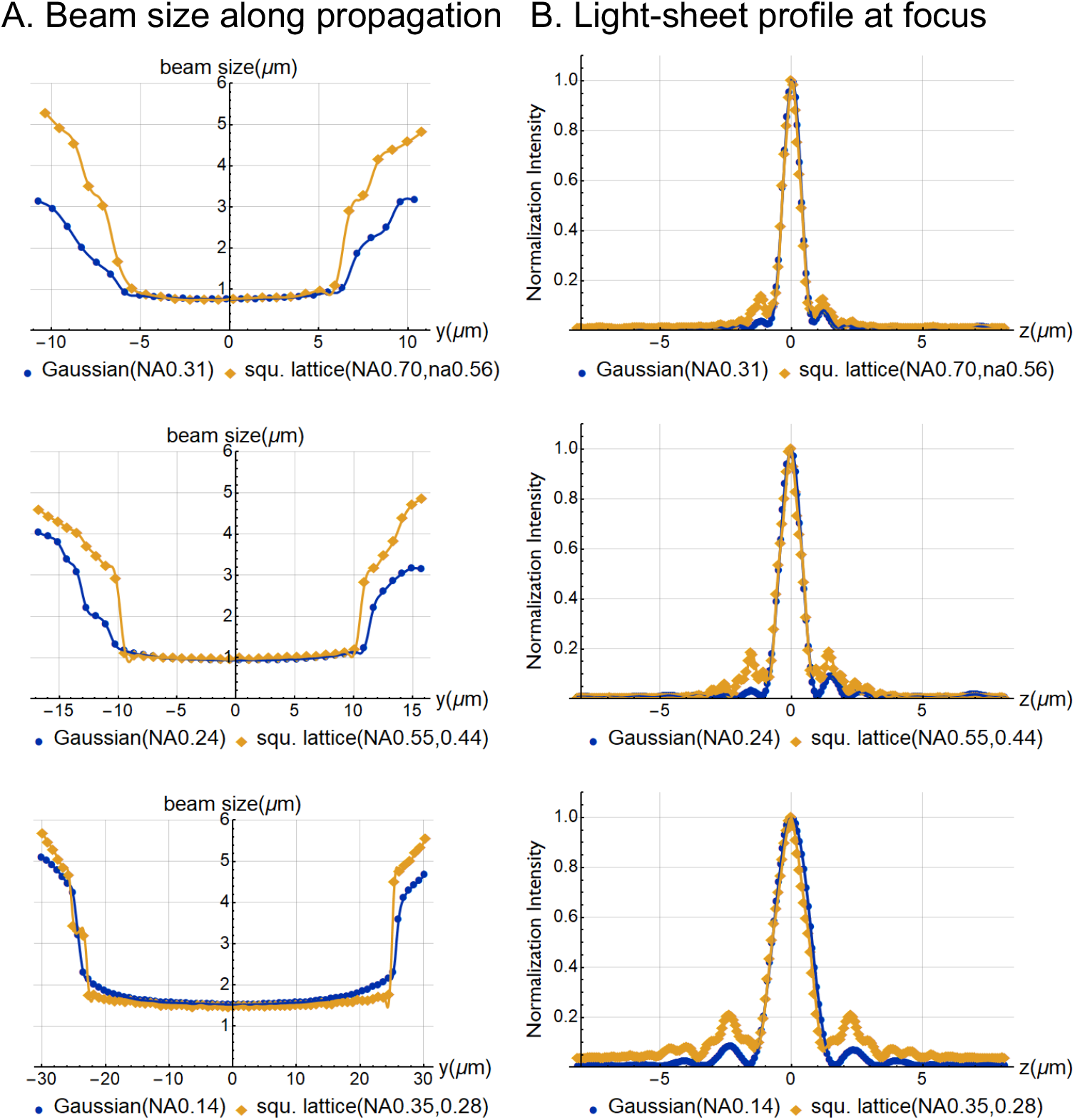
Comparison of light-sheet thickness for three experimentally measured square lattices and Gaussian light-sheets. Gaussian light-sheets were selected for having matching thickness at the beam waist. NA: outer numerical NA of annulus used for squared lattice; na: inner numerical aperture for annulus used for the squared lattice.

A more quantitative analysis of the relationship between light-sheet thickness and propagation length is summarized in Figure 5. Here, each point represents the average of multiple measurements obtained independently from different image sub-volumes of the light-sheet image data, and detailed values of the average and standard deviation are provided in Appendix C. As can be seen, and in agreement with previous numerical simulations [14], the tradeoff between light-sheet propagation length and thickness is similar for both square lattices and Gaussian light-sheets. Of note, the data points for the square lattice show greater variability than those for the Gaussian light-sheets. This arises owing to the larger number of degrees of freedom for generating square lattices. For example, we found that a light-sheet generated from a low NA thick annulus is similar to one generated from a high NA thin annulus (Appendix E). Furthermore, the SLM hologram has parameters that can be tuned, most notably the cropping factor and the spacing, and varying these factors does slightly alter light-sheet properties (Appendix F). While we have used the parameters that have been reported in the lattice literature, one cannot exclude the possibility that through a judicious choice of parameters, more advantageous square lattices could be generated. Nonetheless, in our hands, the best-case lattice light-sheets (i.e. possessing a minimal thickness for a given length) did not outperform the Gaussian light-sheets but followed an almost identical trend for thickness increase with increasing confocal parameter. Furthermore, the data suggest that the inherent complexity of LLSM likely results in greater performance fluctuations than a microscope equipped with a comparatively simple Gaussian light-sheet.

**Fig. 5.**
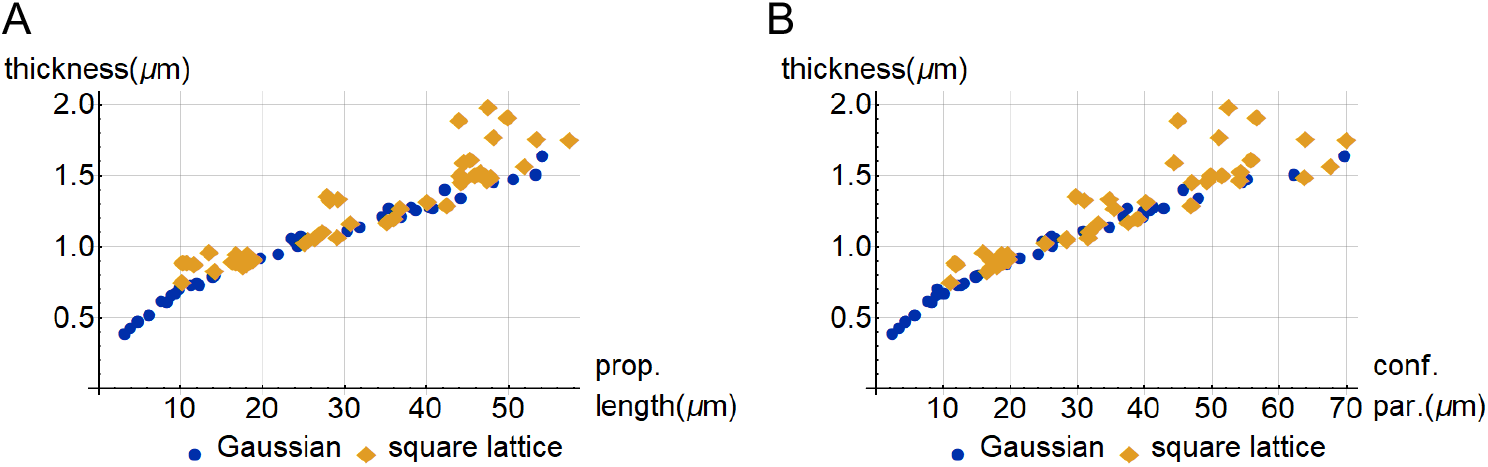
(A) Light-sheet thickness versus light-sheet propagation length. Thickness is measured as described in Figure 2(C) and propagation length is measured as described in Figure 2(D). (B) Light-sheet thickness versus confocal parameter. Confocal parameter is measured as described in Figure 2(E and F).

### 3.2. Comparison of Simulated and Experimentally Measured Light-Sheets

Recently, *Remacha et al* numerically evaluated the optical characteristics for a diverse array of light-sheets (including Gaussian, Bessel, dithered lattice, airy, and more) and concluded that Gaussian beams provided superior image contrast [14]. However, these data were not experimentally confirmed, which is of particular concern for lattice light-sheets since the simulations assumed that each diffraction order in the back focal plane of the illumination objective had constant phase and electric field strength. As such, it has remained an open question if such simulations correctly predicted the properties of experimental lattices. Here we address these concerns and compare our experimental data to theoretical Gaussian beams, and simulated lattice and Gaussian light-sheets. In an effort to maintain consistency, the simulations were performed as described previously [14]. As can be seen in Figure 6, our experimental Gaussian light-sheets are thinner than the theoretical values for Gaussian beams. As mentioned earlier, the intensity distribution in the back focal plane of our illumination objective is more uniform than a traditional Gaussian beam and thus resembles a “flat-top” beam, which could potentially contribute to this discrepancy. Nevertheless, previous simulations by *Remacha et al* demonstrate that Gaussian and “flat-top” beams behave similarly [14], which we performed as well. Indeed, our experimental Gaussian light-sheets are in close correspondence with both the simulated Gaussian and “flat-top” beams, and overall, the difference appears small (Appendix G). Furthermore, both the simulated and experimentally measured “best-case” square lattice light-sheets are in near-perfect agreement, suggesting that the numerical simulations performed by *Remacha et al* were accurate. And lastly, our Gaussian beams are largely indistinguishable from the lattice light-sheets in terms of light-sheet thickness. Importantly, this analysis only takes into account the thickness of the main lobe, while ignoring the sidelobes that are present in the lattice light-sheets.

**Fig. 6.**
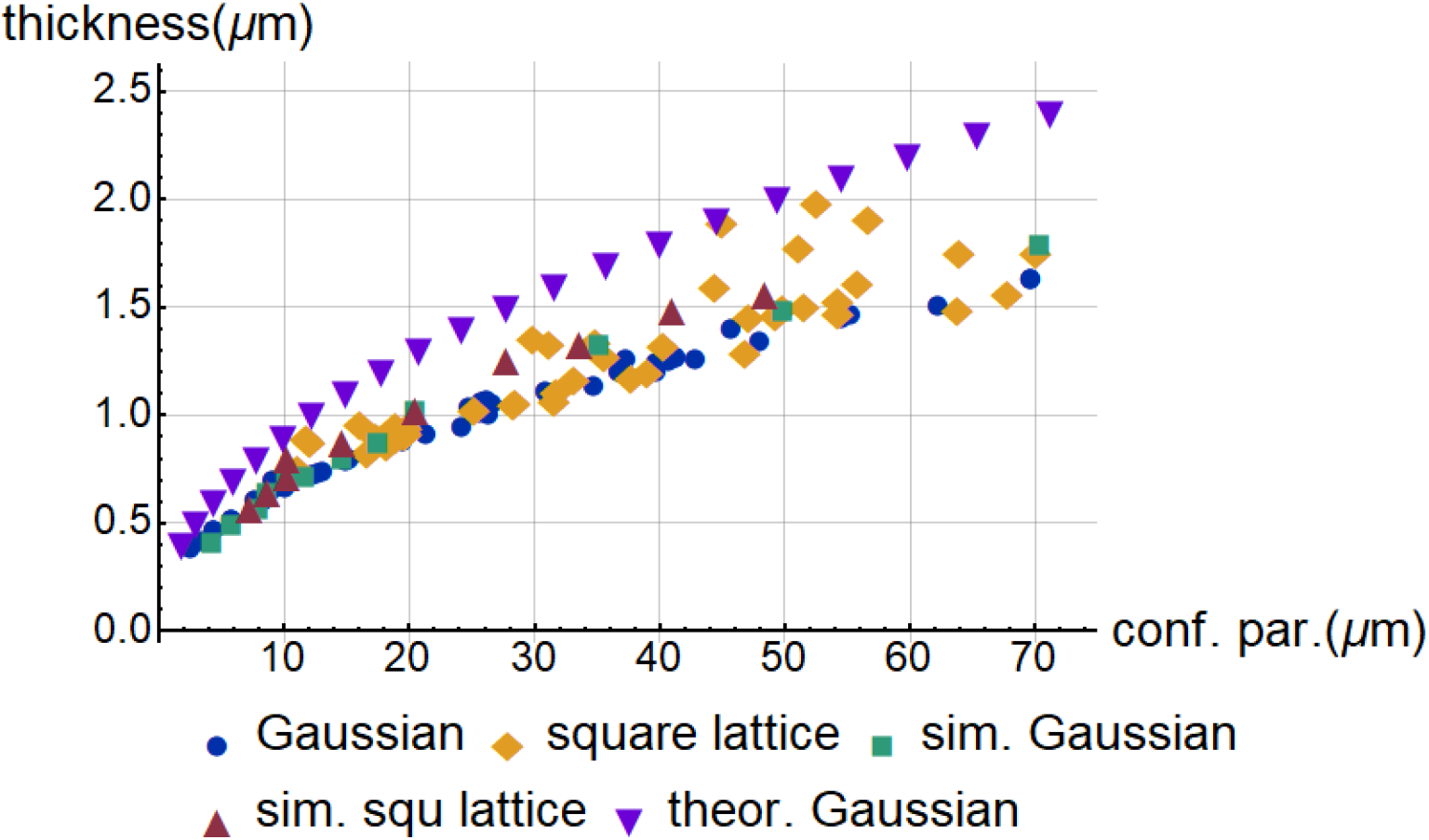
Comparison of experimentally measured Gaussian and square lattice light-sheets with numerically simulated Gaussian, square lattice, and theoretical Gaussian light-sheets derived from an analytical formula.

## 4. Summary

LLSM has generated significant interest in the biological imaging community, with several high-profile publications referring to its illumination optics as “ultrathin” [9–11, 18] and its resolution as “unparalleled” [19]. Yet, as we have shown previously [15], the time-averaged equivalent of these illumination beams can be generated incoherently, and therefore their performance is not a unique consequence of the coherent superposition of multiple Bessel beams. Furthermore, as we show here, the most popular lattice light-sheet, the dithered square lattice, does not appear to provide a significant performance improvement over Gaussian light-sheets, as measured from their beam waist thickness and confocal parameter. We also investigated the Optical Transfer Function for both Gaussian and square lattice light-sheets, and found that both Optical Transfer Functions extend to a similar level (Appendix H). Thus, for LSFM, these beams would result in practically indistinguishable levels of optical sectioning, field of view, and axial resolution.

According to both numerical simulations [8, 14] and the experimental measurements, the square lattice does not behave like a propagation invariant beam, but rather a divergent one. Indeed, its tendency to increase in thickness as the confocal parameter grows is in close agreement to a simple Gaussian light-sheet. This in itself is a surprise, as square optical lattices are produced by a coherent superposition of Bessel beams, which are *de facto* propagation invariant. Therefore, we conclude that for certain periodicities of the lattice (i.e. corresponding to the square lattice), the propagation invariant nature of the parent beam can be lost, and a clear explanation for this phenomenon is still under investigation.

For higher resolution imaging, the coherent modulation present in static (e.g., non-dithered) lattice light-sheets can be used for structured illumination microscopy [9]. However, owing to the LSFM geometry, this mode improves the resolution only in the axial dimension and one of the two lateral dimensions (e.g., the X-direction). As such, we believe that the main benefit of optical lattices for structured illumination microscopy is that they provide greater frequency support in the axial dimension which cannot otherwise be achieved yet with one dimensional beam engineering. Nonetheless, owing to the need to acquire 5 images for one structured illumination reconstruction, and the fact that the lateral resolution gain is modest, the use of this LLSM mode has so far been rare. Alternatively, hexagonal lattices provide a thinner main lobe than square lattices at the cost of much stronger sidelobes that grow drastically with the light-sheet propagation length. Indeed, sidelobes that approach 50% strength of the main lobe are notoriously hard to remove computationally [20], which is likely the main reason why hexagonal lattice light-sheets have seen little use [21]. And lastly, it appears to us that both for the square and hexagonal lattices, the reported resolutions could only have been achieved via strong deconvolution of the data.

In conclusion, as the vast majority of LLSM is performed with the dithered square lattice and not the structured illumination or hexagonal lattice light-sheet modes, our results stipulate that this type of imaging could alternatively be performed with conventional Gaussian or flat-top light-sheets. This would be a welcome simplification of such high-resolution light-sheet microscopes, as the correct alignment of a LLSM setup requires optical expertise. Furthermore, by dispensing with the spatial light-modulator, much less input laser power is needed, and simultaneous multicolor excitation is also easily achieved. Importantly, we interpret our results as a poignant reminder that there is no free lunch in linear optics; with increasing confocal parameter, it appears that one either has to accept a thicker light-sheet, stronger sidelobes, or both. Lattice light-sheets have been heralded as a breakthrough that overcomes this trade-off. Nonetheless, our results indicate that they underlie the same physics and limitations as a simple Gaussian light-sheet.

## 5.2 Acknowledgments

The authors would like to thank Florian Fahrbach for help with the simulation code and critical reading of the manuscript. Furthermore, we are grateful to Dr. Kim Reed for her continued support, as well as all the employees at the University of Texas Southwestern Medical Center who make our research possible.

## 7. Article thumbnail upload

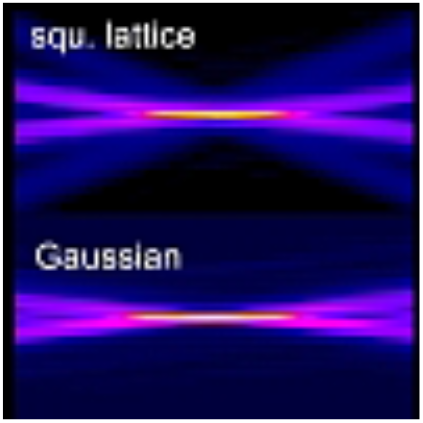

## Appendix A. Experimental System

**Fig. S1.**
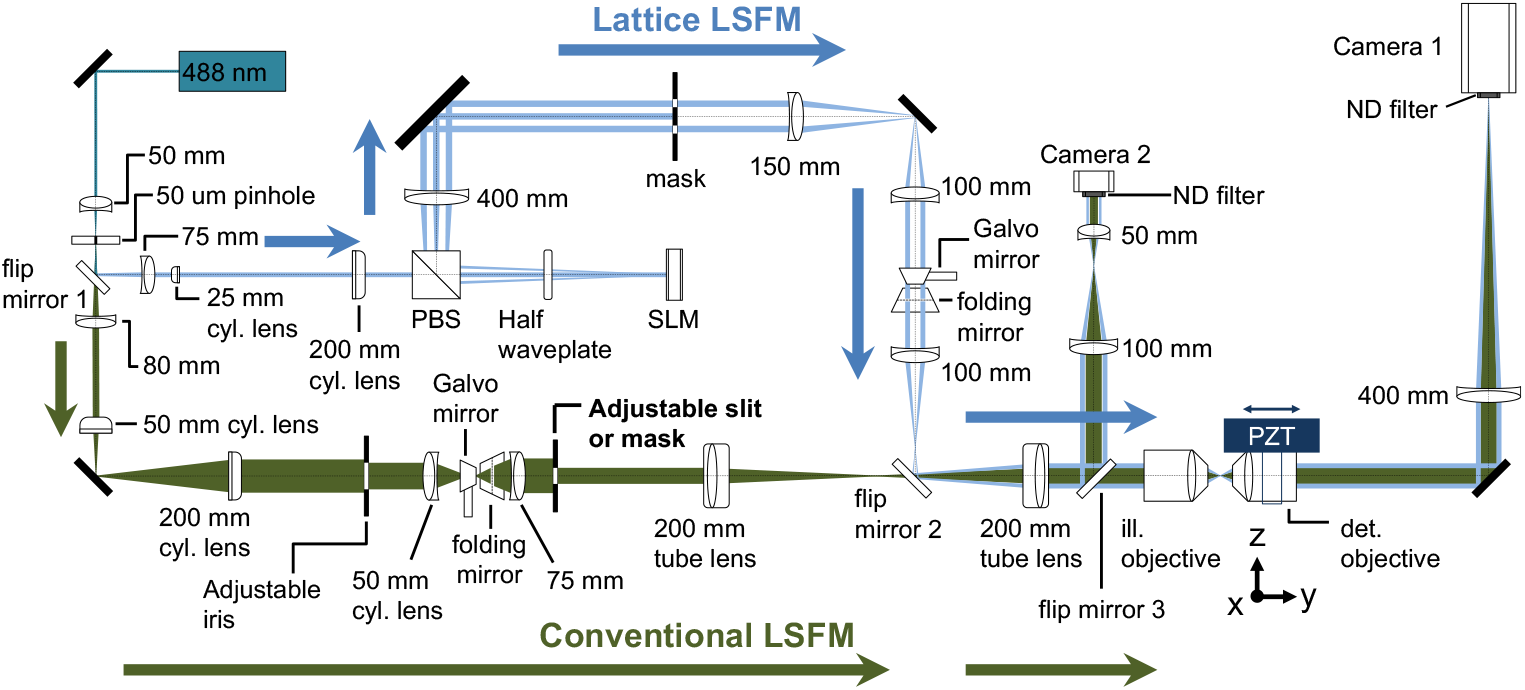
Schematic representation of the experimental setup, which consists of two illumination paths, one to produce optical lattices (blue arrow “Lattice LSFM”) and one to produce conventional Gaussian light-sheets (green arrow “Conventional LSFM”). Both light-sheets are ultimately injected into the illumination objective (ill. objective). A secondary objective (det. objective) observes the image of the light-sheets in transmission and projects them onto a scientific CMOS camera (Camera 1). Flip mirror 1 and 2 are used to select between the Lattice LSFM or the conventional LSFM light path. An image of the back focal plane of the illumination objective can be observed on Camera 2 by directing the laser light with flip mirror 3. SLM: Spatial light modulator; ND: neutral density filter; cyl. lens: cylindrical lens.

## Appendix B. Effective NA of Gaussian light-sheet generated with an annulus

**Fig. S2.**
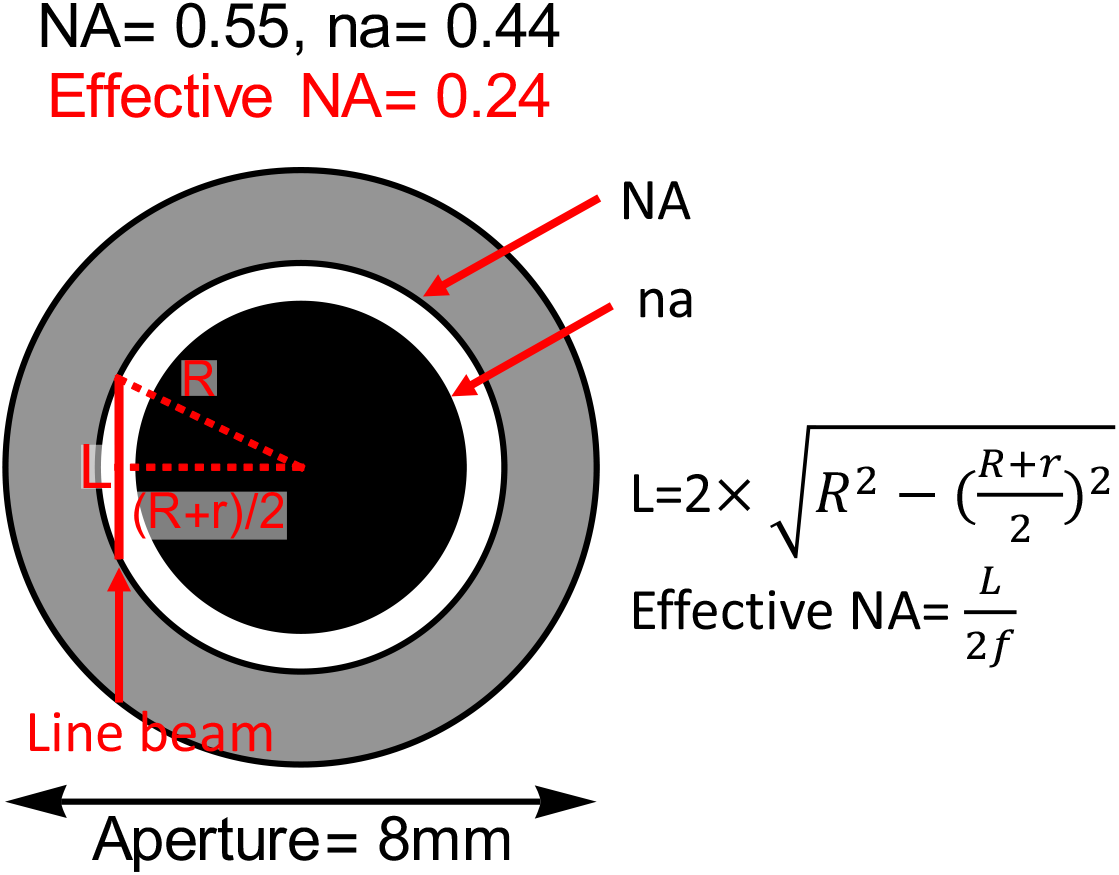
While we primarily used an adjustable slit to create the different Gaussian light-sheets, some datasets were acquired using an annular mask, and placing the line focus at the periphery of the annulus. To generate a 1D Gaussian light-sheet, a subset of the back pupil of the objective is illuminated with a line focus, and the length of the line defines its effective NA. The effective NA can be calculated as the equation shown here. For example, the back-pupil of a Nikon 40x NA0.8 objective is 8 mm in diameter. By using an annular mask with NA=0.55 and na=0.44 results in an effective NA of 0.24.

## Appendix C. Systematic Comparison of Gaussian and square lattice Light-Sheets

Table S1 summarizes all Gaussian and square lattice light-sheets that we have measured. In total 39 and 52 experiments have been performed for Gaussian and square lattice light-sheets, respectively. The data covers a wide range to illustrate the characteristics of both Gaussian and square lattice light-sheets. In the square lattice, NA and na represent the outer and inner numerical apertures of the annulus, respectively. Bold indicates that the light-sheets were used in the original Lattice Light-Sheet Microscopy manuscript [14]. For Gaussian beams, the effective NA used to generate the light-sheet was controlled in most cases with a micrometer slit that is conjugate to the back-pupil of the illumination objective. In some cases, we replace the micrometer slit with an annulus. In such case, the annulus served as the slit, and the effective NA is reported as described in Figure S2.

**Table S1.** Summary of all experimental data shown in Figure 5. The table is sorted with thickness and then the confocal parameter. Eff. NA represents the effective NA used for Gaussian beams, which corresponds to the NA of the slit or the theoretical NA of the annulus. In the square lattice, sp and cp represents spacing and cropping factor, respectively. n represents the number of measurements. For Gaussian measurements, n equals to 24 for all measurements. In the square lattice measurements, n equals 6 unless it is specified otherwise.

| Gaussian |  |  |  | Square lattice |  |  |  |
| --- | --- | --- | --- | --- | --- | --- | --- |
| Thickness | Propagation length | Confocal parameter | eff. NA | Thickness | Propagation length | Confocal parameter | NA,na (sp.,cp., (n)) |
| 0.4±0.01 | 3.3±0.09 | 2.6±0.04 | 0.61 | 0.8±0.02 | 10.3±0.16 | 11.2±0.20 | 0.70,0.56 |
| 0.4±0.00 | 4.0±0.15 | 3.5±0.08 | 0.54 | 0.8±0.01 | 14.3±0.11 | 16.7±0.24 | 0.70,0.60 |
| 0.5±0.01 | 4.9±0.27 | 4.5±0.14 | 0.49 | 0.9±0.04 | 10.9±0.30 | 11.9±0.46 | 0.70,0.51 |
| 0.5±0.00 | 6.3±0.26 | 5.9±0.28 | 0.42 | 0.9±0.05 | 11.8±0.44 | 12.1±0.68 | 0.70,0.56 |
| 0.6±0.01 | 7.7±0.24 | 7.8±0.31 | 0.35 | 0.9±0.02 | 17.7±0.52 | 17.5±1.41 | **0.55,0.44** |
| 0.6±0.01 | 8.5±0.12 | 8.4±0.25 | 0.36 | 0.9±0.00 | 16.5±0.15 | 17.9±0.44 | **0.55,0.44** |
| 0.7±0.02 | 9.0±0.38 | 9.2±0.10 | 0.32 | 0.9±0.01 | 18.2±0.17 | 17.9±0.35 | **0.55,0.44** |
| 0.7±0.02 | 9.9±0.40 | 9.2±0.88 | 0.35 | 0.9±0.02 | 17.7±0.64 | 18.2±1.07 | **0.55,0.44** |
| 0.7±0.02 | 9.5±0.34 | 10.2±0.69 | 0.32 | 0.9±0.01 | 18.2±0.15 | 18.3±0.32 | **0.55,0.44** |
| 0.7±0.01 | 11.4±0.45 | 12.3±0.27 | 0.28 | 0.9±0.01 | 10.4±0.10 | 18.5±0.34 | **0.55,0.44** |
| 0.7±0.01 | 12.4±0.08 | 12.7±0.23 | 0.31 | 0.9±0.01 | 16.9±0.18 | 18.6±0.26 | 0.55,0.40 |
| 0.7±0.04 | 12.0±0.78 | 13.2±0.98 | 0.28 | 0.9±0.01 | 17.9±0.28 | 18.6±0.33 | **0.55,0.44** |
| 0.8±0.02 | 14.1±0.61 | 15.0±0.77 | 0.26 | 0.9±0.02 | 18.1±0.65 | 18.6±0.81 | 0.55,0.52 |
| 0.8±0.02 | 14.4±0.77 | 15.3±1.21 | 0.26 | 0.9±0.01 | 17.5±0.19 | 18.7±0.38 | **0.55,0.44** |
| 0.9±0.02 | 18.0±0.93 | 19.5±0.73 | 0.22 | 0.9±0.01 | 17.3±0.24 | 19.0±0.68 | **0.55,0.44** |
| 0.9±0.02 | 19.8±0.27 | 21.4±0.24 | 0.24 | 0.9±0.01 | 19.0±0.46 | 19.2±0.91 | **0.55,0.44** |
| 0.9±0.03 | 22.0±1.46 | 24.2±1.59 | 0.22 | 0.9±0.02 | 17.9±0.23 | 19.8±0.67 | **0.55,0.44** |
| 1.0±0.03 | 23.9±1.20 | 24.9±1.01 | 0.20 | 0.9±0.01 | 18.0±0.12 | 19.8±0.05 | **0.55,0.44** |
| 1.0±0.03 | 24.3±1.90 | 25.7±1.87 | 0.20 | 0.9±0.01 | 18.3±0.26 | 19.8±0.65 | **0.55,0.44** |
| 1.0±0.03 | 24.4±2.00 | 26.4±2.41 | 0.20 | 0.9±0.02 | 18.9±0.23 | 19.8±0.45 | **0.55,0.44** |
| 1.1±0.02 | 23.6±1.17 | 25.9±1.11 | 0.20 | 1.0±0.04 | 13.6±0.21 | 16.1±0.26 | 0.70,0.60 |
| 1.1±0.02 | 24.7±1.61 | 26.3±2.27 | 0.20 | 1.0±0.01 | 17.0±0.24 | 19.0±0.32 | 0.55,0.40 |
| 1.1±0.04 | 25.2±1.96 | 26.7±1.46 | 0.20 | 1.0±0.01 | 25.3±0.61 | 25.3±0.82 | **0.51,0.43 (n=18)** |
| 1.1±0.03 | 30.5±1.76 | 31.0±0.74 | 0.18 | 1.0±0.01 | 25.7±0.41 | 28.5±0.61 | **0.51,0.43** |
| 1.1±0.05 | 32.0±1.73 | 34.8±2.06 | 0.18 | 1.1±0.02 | 26.5±0.23 | 28.5±0.77 | **0.51,0.43** |
| 1.2±0.02 | 34.7±0.83 | 36.9±2.41 | 0.16 | 1.1±0.01 | 29.3±0.30 | 31.7±0.57 | **0.55,0.48**<br>(n=12) |
| 1.2±0.02 | 37.0±0.4 | 39.8±0.92 | 0.18 | 1.1±0.01 | 27.5±0.34 | 31.9±0.60 | **0.55,0.48** |
| 1.3±0.09 | 35.4±2.79 | 37.5±1.17 | 0.16 | 1.2±0.01 | 30.9±1.36 | 33.2±0.22 | **0.55,0.48** |
| 1.3±0.04 | 36.8±1.89 | 39.8±2.16 | 0.16 | 1.2±0.02 | 35.3±0.42 | 37.8±1.27 | 0.40,0.32 |
| 1.3±0.05 | 38.8±2.36 | 40.8±3.35 | 0.16 | 1.2±0.01 | 36.1±0.85 | 39.1±1.01 | 0.40,0.32<br>(n=18) |
| 1.3±0.02 | 38.2±0.86 | 41.4±2.36 | 0.16 | 1.3±0.02 | 28.4±0.35 | 31.2±0.52 | 0.70,0.67 |
| 1.3±0.02 | 40.4±0.75 | 41.4±2.43 | 0.16 | 1.3±0.01 | 29.4±0.44 | 34.9±0.93 | 0.70,0.67 |
| 1.3±0.03 | 40.8±1.64 | 42.9±2.43 | 0.16 | 1.3±0.02 | 36.9±1.05 | 35.6±1.35 | 0.55,0.52 |
| 1.3±0.06 | 44.2±3.36 | 48.0±0.81 | 0.14 | 1.3±0.02 | 40.2±0.86 | 40.4±0.99 | 0.55,0.52<br>(sp0.93,<br>cp0.05) |
| 1.4±0.09 | 42.3±4.55 | 45.8±1.56 | 0.16 | 1.3±0.01 | 42.6±0.48 | 47.0±0.84 | 0.35,0.28 |
| 1.5±0.08 | 48.2±2.55 | 54.7±3.34 | 0.14 | 1.4±0.02 | 28.0±0.79 | 30.0±1.19 | 0.70,0.67 |
| 1.5±0.03 | 50.6±1.66 | 55.4±1.30 | 0.14 | 1.5±0.02 | 44.3±0.85 | 47.2±1.67 | 0.55,0.52<br>(sp0.93,<br>cp0.10) |
| 1.5±0.04 | 53.4±1.53 | 62.4±4.05 | 0.14 | 1.5±0.01 | 44.5±2.46 | 49.4±1.99 | 0.55,0.52 |
| 1.6±0.03 | 54.2±2.67 | 69.8±3.80 | 0.13 | 1.5±0.02 | 44.2±1.16 | 49.9±0.44 | 0.55,0.52 |
|  |  |  |  | 1.5±0.03 | 46.0±0.71 | 51.6±0.73 | 0.55,0.52 |
|  |  |  |  | 1.5±0.02 | 46.7±0.63 | 54.4±2.19 | 0.55,0.52<br>(sp0.93,<br>cp0.15) |
|  |  |  |  | 1.5±0.08 | 47.5±2.43 | 54.4±3.10 | 0.55,0.52 |
|  |  |  |  | 1.5±0.02 | 48.0±2.38 | 63.8±2.20 | 0.55,0.52<br>(sp0.92,<br>cp0.05) |
|  |  |  |  | 1.6±0.02 | 44.7±1.17 | 44.5±1.25 | 0.55,0.52<br>(sp0.93,<br>cp0.25) |
|  |  |  |  | 1.6±0.16 | 45.5±2.09 | 56.0±11.75 | 0.55,0.52<br>(sp0.93,<br>cp0.15) |
|  |  |  |  | 1.6±0.11 | 52.1±2.35 | 67.8±8.24 | 0.55,0.52 |
|  |  |  |  | 1.8±0.13 | 48.3±1.59 | 51.2±7.98 | 0.55,0.52<br>(sp0.92,<br>cp0.15) |
|  |  |  |  | 1.8±0.02 | 53.5±1.22 | 64.1±2.33 | 0.55,0.52(<br>sp0.92,<br>cp0.15) |
|  |  |  |  | 1.8±0.14 | 57.5±1.74 | 70.2±7.37 | 0.55,0.52<br>(sp0.91,<br>cp0.15) |
|  |  |  |  | 1.9±0.16 | 44.1±4.31 | 45.1±6.44 | 0.55,0.52<br>(sp0.92,<br>cp0.25) |
|  |  |  |  | 1.9±0.16 | 50.0±3.20 | 56.8±8.31 | 0.55,0.52(<br>sp0.91,<br>cp0.25) |
| | | | | $2.0 \pm 0.22$ | $47.6 \pm 1.97$ | $52.6 \pm 6.36$ | $0.55, 0.52$<br>(sp0.93,<br>cp0.25) |

## Appendix D. Contribution of Zeroth Order Beams to Square Lattice Properties

We performed measurements on square lattices before and after blocking the zeroth, +1, and −1 diffraction orders. Only one data set is shown for each annulus, however similar results can be found with other measurements. Table S2 summarizes the measurements with multiple annuli. It can also be seen that when the 0th order is blocked or only +1 or −1 is allowed, the light-sheet still shows similar properties to the original dithered square lattice light-sheet

**Table S2.** Dependence of Square Lattice Properties on Back-Pupil Diffraction Orders

|  |  |  |  |
| --- | --- | --- | --- |
| NA 0.35, na 0.28 | Dithered squ. lattice | 0 order blocked | Only +1 or -1 order |
| Thickness ($\mu\text{m}$) | 1.3± 0.01 | 1.3± 0.01 | |
| Axial FWHM ($\mu\text{m}$) | 42.6± 0.48 | 43.1± 0.47 | |
| Confocal parameter ($\mu\text{m}$) | 47.0± 0.84 | 50.1± 2.43 | |
| NA 0.55, na 0.40 | Dithered squ. lattice | 0 order blocked | Only +1 or -1 order |
| Thickness ($\mu\text{m}$) | 0.9± 0.01 | 0.9± 0.01 | |
| Axial FWHM ($\mu\text{m}$) | 16.9± 0.18 | 17.5± 0.63 | |
| Confocal parameter ($\mu\text{m}$) | 18.6± 0.26 | 17.2± 0.60 | |
| NA 0.55, na 0.44 | Dithered squ. lattice | 0 order blocked | Only +1 or -1 order |
| Thickness ($\mu\text{m}$) | 0.9± 0.02 | 0.8± 0.01 | 0.8± 0.01 |
| Axial FWHM ($\mu\text{m}$) | 17.3± 0.24 | 18.1± 0.21 | 18.3± 0.75 |
| Confocal parameter ($\mu\text{m}$) | 19.0± 0.68 | 18.5± 0.62 | 18.8± 0.89 |
| NA 0.55, na 0.48 | Dithered squ. lattice | 0 order blocked | Only +1 or -1 order |
| Thickness ($\mu\text{m}$) | 1.2± 0.01 | 1.2± 0.01 | |
| Axial FWHM ($\mu\text{m}$) | 30.9± 1.36 | 26.9± 0.90 | |
| Confocal parameter ($\mu\text{m}$) | 33.2± 0.22 | 32.2± 0.37 | |
| NA 0.55, na 0.52 | Dithered squ. lattice | 0 order blocked | Only +1 or -1 order |
| Thickness ($\mu\text{m}$) | 1.5± 0.01 | 1.5± 0.01 | |
| Axial FWHM ($\mu\text{m}$) | 44.5± 2.46 | 44.2± 2.45 | |
| Confocal parameter ($\mu\text{m}$) | 49.4± 1.99 | 50.0± 2.56 | |
| NA 0.70, na 0.56 | Dithered squ. lattice | 0 order blocked | Only +1 or -1 order |
| Thickness ($\mu\text{m}$) | 0.8± 0.02 | 0.7± 0.02 | |
| Axial FWHM ($\mu\text{m}$) | 10.3± 0.16 | 10.8± 0.14 | |
| Confocal parameter ($\mu\text{m}$) | 11.2± 0.20 | 11.1± 0.25 | |
| NA 0.70, na 0.60 | Dithered squ. lattice | 0 order blocked | Only +1 or -1 order |
| Thickness ($\mu\text{m}$) | 0.8± 0.01 | 0.8± 0.01 | |
| Axial FWHM ($\mu\text{m}$) | 14.3± 0.11 | 14.7± 0.28 | |
| Confocal parameter ($\mu\text{m}$) | 16.7± 0.24 | 16.9± 0.23 | |
| NA 0.70, na 0.67 | Dithered squ. lattice | 0 order blocked | Only +1 or -1 order |
| Thickness ($\mu\text{m}$) | 1.3± 0.01 | 1.4± 0.03 | |
| Axial FWHM ($\mu\text{m}$) | 29.4± 0.44 | 30.7± 3.01 | |
| Confocal parameter ($\mu\text{m}$) | 34.9± 0.93 | 37.0± 1.80 | |

**Fig. S3.**
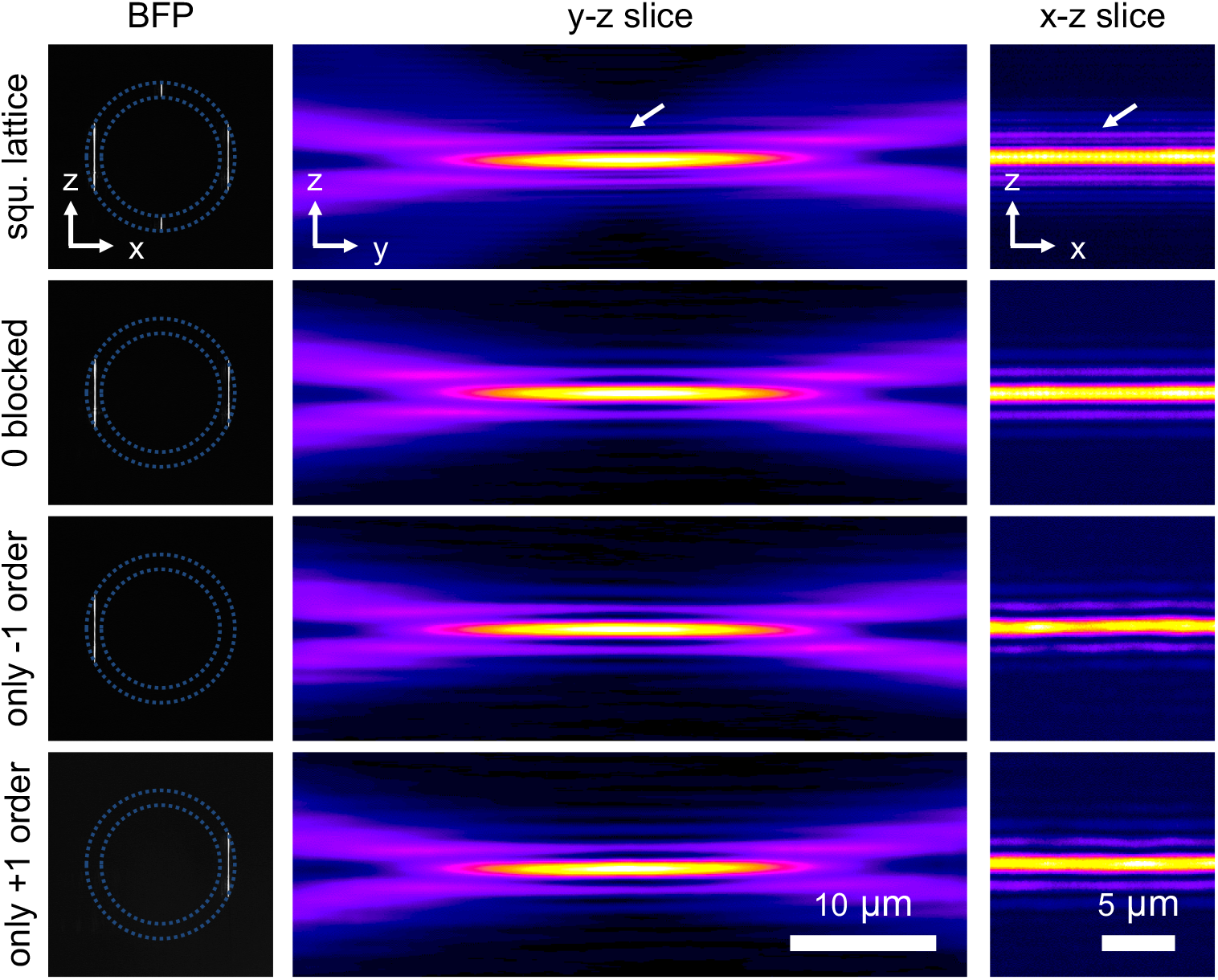
Light-sheet generated by blocking 0 order or allowing only +1 or −1 order arising from the phase hologram used in the lattice LSFM setup. Images of the back focal plane (BFP) of the illumination objective, and YZ, and XZ slices of the light-sheets are shown. Sidelobes are present in all conditions, except for the one with all orders present (square lattice) they appear more prominent. The annulus used here possesses and outer and inner NA of 0.55 and 0.44, respectively.

## Appendix E. Choice of Annulus for Lattice Generation

As shown in Figure S4(A), we found similar light-sheets can be generated using different annuli. Interestingly, the thickness and the confocal parameter are 0.9±0.01 μm and 18.6±0.26 μm from an NA=0.55, na=0.40 annulus, respectively; and they are 1.0±0.04 μm and 16.1±0.26 μm from an NA=0.70, na=0.60 annulus, respectively. Thus, the square lattice light-sheet generated with the lower NA annulus is more desirable (thinner and longer) in this case.

This can be explained in Figure S4(B), where the square lattice light-sheet can be considered as the interference of four light-sheets. These four light-sheets can be grouped into two pairs: the “long pair” corresponds to the two long line beams in the back focal plane and they are displaced in k_x_, and the “short pair” corresponds to the two short line beams that are and displaced in k_z_ (k being the coordinate in reciprocal space). The long pair dominates the thickness and the length of the final lattice light-sheet, whereas the short pair may reduce the thickness but also shorten the length. As shown in the first panel of Figure S4(C), L and the corresponding effective NA can be calculated from the outer and inner radii of the annulus. In the example shown here, the effective NA of NA=0.55, na=0.40, and NA=0.70, na=0.60 are 0.28 and 0.26, respectively. Therefore, we can expect the lattice light-sheet generated from NA=0.55, na=0.40 would be thinner and it is also supported by our result. The lattice light-sheet generated form NA=0.70, 0.60 annulus is indeed slightly thicker, but surprisingly, it is also shorter than the one generated from NA0.55, na0.40 annulus. This could be due to fact that the short pair in the NA=0.70, 0.60 annulus contributes less in the improvement of the thickness, but they come from a steeper angle in Z, thus making the light-sheet shorter.

In short, to generate a better square lattice light-sheet with a high NA annulus probably needs more careful adjustment. This is because the high NA annulus usually gives lower effective NA. To make it thinner the short pair need to be tuned carefully but it is also constrained by the annulus, therefore a special mask that is not shaped like an annulus might be needed.

**Fig. S4.**
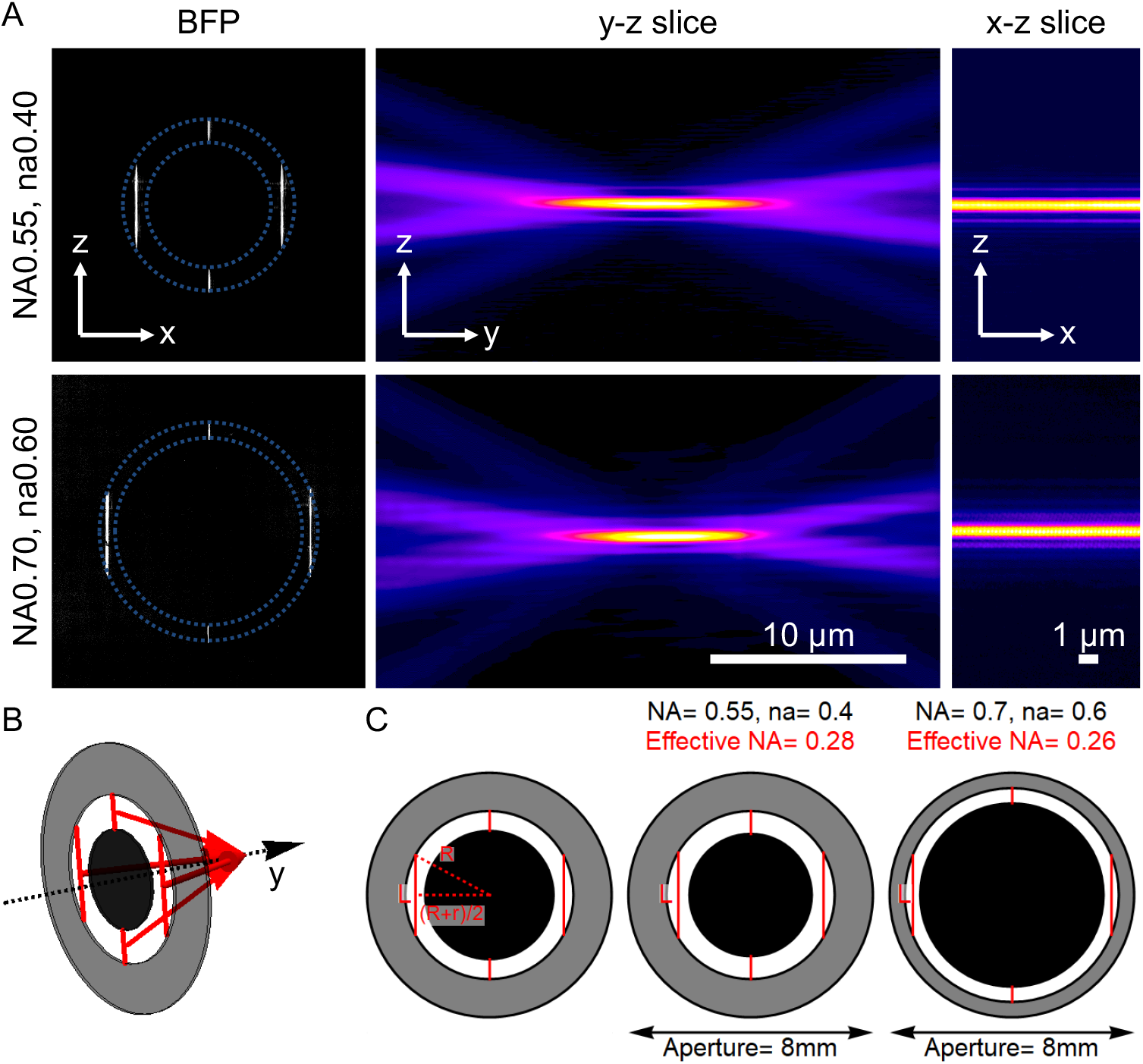
Comparison of similar square lattice light-sheet generated with different annuli. (A) Images of the back focal plane of the illumination objective, YZ slices, and XZ slices of the light-sheets. (B) The schematic sketch illustrates that the square lattice light-sheet is the result of the interference of four light-sheets. (C) From left to right: the calculation of effective NA, the effective NA of the NA0.55, na0.44 and NA0.7, na0.60 annulus. The effective NA is obtained from 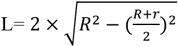.

## Appendix F. Choice of SLM Parameters for Lattice Generation

The SLM phase hologram is essential for generating lattice light-sheets. The code we are using to generate the SLM phase hologram basically follows the practical optical path. It first generates the pattern on the back focal plane (BFP) of the illumination objective by considering that the lattice pattern is the interference of a group of Bessel beams. The Fourier transform of the pattern on the BFP of the illumination objective is then the basis of the pattern put on the SLM. The SLM pattern is affected by the size of the annulus, the wavelength, and the magnification of the optical path. Nevertheless, to obtain a proper lattice, several parameters need to be adjusted carefully, and the cropping factor and the spacing are the two main parameters.

The cropping factor is basically a threshold that crops some residues in the SLM phase hologram. In practice, it affects the sidelobes in the final lattice light-sheet. As shown in Figure S5 indicated with the white arrows, with larger cropping factor, the sidelobes and the intensity of the short line-beams are decreasing, and the lattice becomes more Gaussian like. Please also note that in Figure S5, spacing = 0.94, large sidelobes appear. So even with very large cropping factor it does not resemble anymore to a Gaussian light-sheet. Nevertheless, we can still see the side-lobes are decreasing with larger cropping factor.

The spacing parameter basically determines the period of the lateral periodic pattern of the lattice. Effectively, it also determines the position of the long line-beams, or the distance between them, as indicated by the red dashed line in Figure S5. Generally, the spacing parameter is chosen when the long line-beams are in the middle of the outer and inner rings of the annulus. However, there is still some flexibility in their fine spacing. Basically, the larger the spacing parameter, the closer the two line-beams, and the light-sheet is also shorter because the long line-beams are getting longer, as explained in Figure S4(B and C). Please also note that for a spacing = 0.94, strong sidelobes that are characteristic for hexagonal lattice patterns start to appear.

Table S3 shows the quantifications of the light-sheets shown in Figure S5, which also supports the aforementioned observation. We also put two Gaussian light-sheet that has similar thickness for comparison, and they show similar properties to the dithered square lattice light-sheet. We can see that with the same thickness (~1.5 μm), the Gaussian light-sheet has a length (~62-64 μm) that is very similar to the longest dithered square lattice (spacing = 0.92, cropping factor = 0.05) in Figure S5. The light-sheet length increases (69.8 μm) along with an increase of its thickness. We did not include the dithered square lattice light-sheet with spacing= 0.94 because it behaves more like a hexagonal lattice light-sheet and the quantification of it is very different.

**Table S3.** The thickness and confocal parameters for dithered square lattice light-sheets using different spacing and cropping factors are shown. The numbers for the lattice light-sheet with spacing= 0.94 are not shown because it already does not behave like a square lattice. The numbers of two Gaussian light-sheets that have similar properties are shown for comparison. n=6 for the lattice light-sheet measurements and n= 24 for the Gaussian light-sheet measurements have been used to compute average and standard deviation of the different parameters. cp= cropping factor.

|  | Dithered Square lattice light-sheet |  |  |  |  | Gaussian |  |
| --- | --- | --- | --- | --- | --- | --- | --- |
|  | Spacing= 0.92 |  | Spacing= 0.93 |  |  | Effective NA = 0.142 | Effective NA = 0.128 |
|  | cp=0.05 | cp=0.15 | cp=0.05 | cp=0.10 | cp=0.15 |  |  |
| **Thickness ($\mu\text{m}$)** | 1.5<br>$\pm 0.02$ | 1.8<br>$\pm 0.02$ | 1.3<br>$\pm 0.02$ | 1.5<br>$\pm 0.02$ | 1.5<br>$\pm 0.02$ | 1.5<br>$\pm 0.04$ | 1.6<br>$\pm 0.03$ |
| **Propagation length ($\mu\text{m}$)** | 48.0<br>$\pm 2.38$ | 53.5<br>$\pm 1.22$ | 40.2<br>$\pm 0.86$ | 44.3<br>$\pm 0.85$ | 46.7<br>$\pm 0.63$ | 53.4<br>$\pm 1.53$ | 53.9<br>$\pm 2.85$ |
| **Confocal Parameter ($\mu\text{m}$)** | 63.8<br>$\pm 2.20$ | 64.1<br>$\pm 2.33$ | 40.4<br>$\pm 0.99$ | 47.2<br>$\pm 1.67$ | 54.4<br>$\pm 2.19$ | 62.4<br>$\pm 4.05$ | 69.8<br>$\pm 3.72$ |

**Fig. S5.**
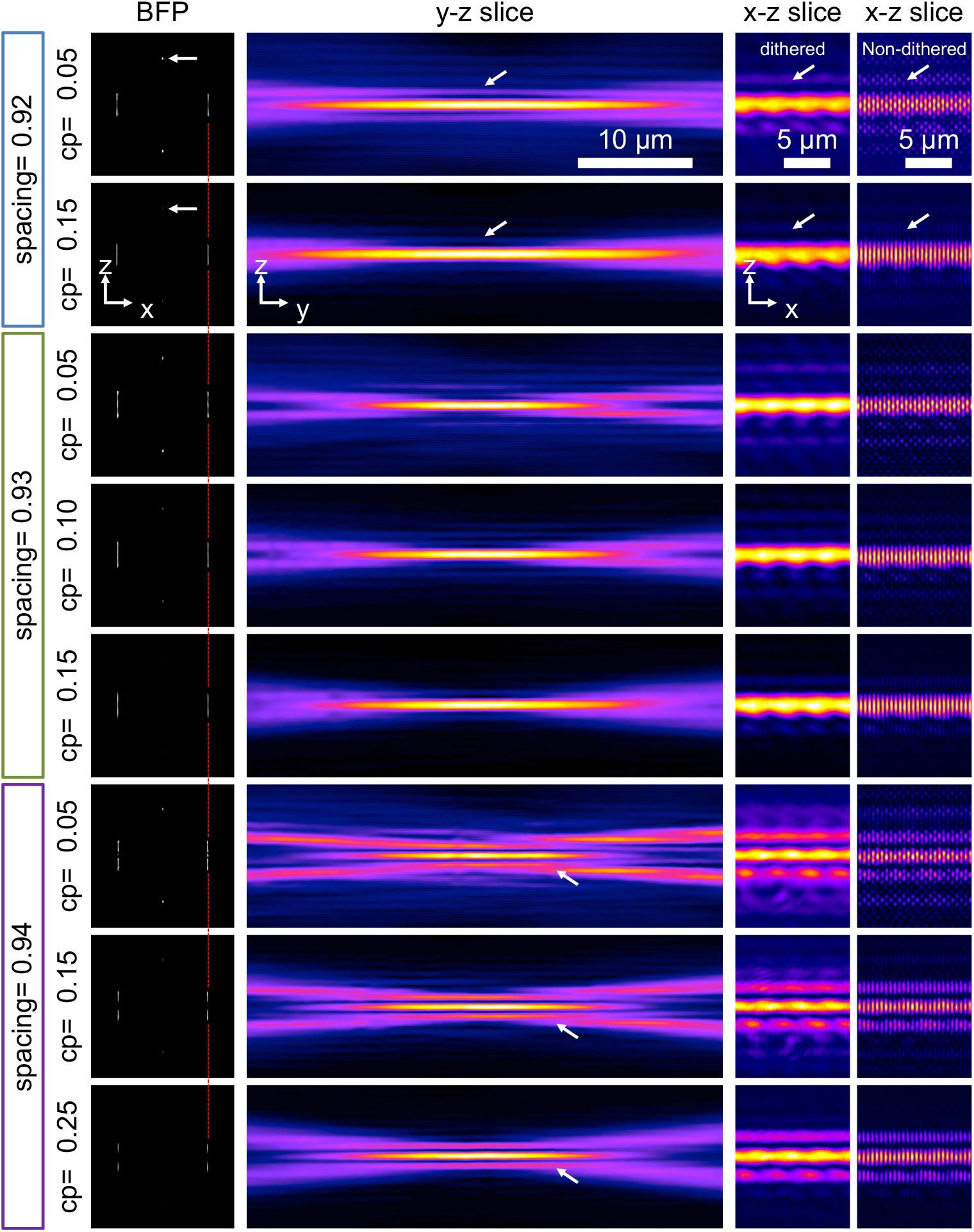
Effect of cropping factor and spacing on lattice light-sheet generation. The combination of three spacing numbers and different cropping factors are shown. From left to right, the line beams on the Back focal plane (BFP) of the illumination objective, YZ and XZ slices are shown. The XZ slice of the non-dithered lattice is shown in the rightmost column. Note that the scaling in the XZ slice is different from the YZ in order to show clearer the lattice pattern and the side lobes. In the BFP, white arrows point to the short line beams and how they are affected by the cropping factor (cp). The red dashed line indicates the shift of the long line beams depending on the spacing. In the YZ and XZ slices, white arrows point to the appearance and disappearance of the sidelobes. The NA and na of the annulus used in this example are 0.55 and 0.52, respectively.

## Appendix G. Gaussian versus flat top light-sheets

**Fig. S6.**
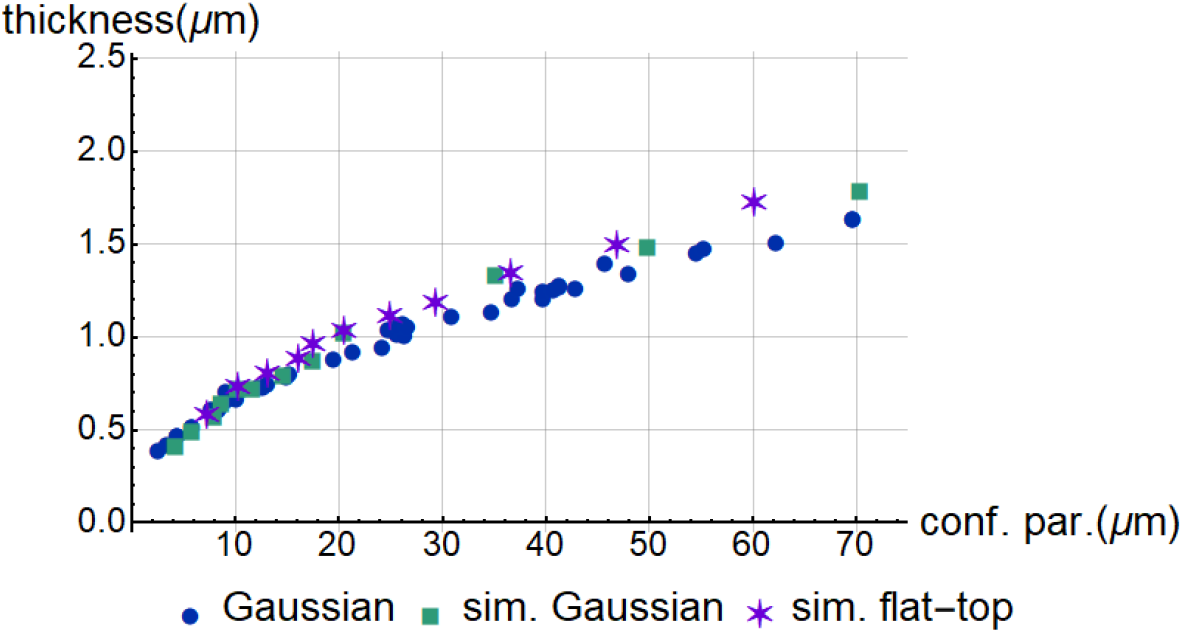
Our experimentally measured Gaussian light-sheets compared to simulated Gaussian and flat-top light-sheets. A flat-top light-sheet has a constant intensity profile in the back focal plane. Here we can see the characteristics of simulated Gaussian and flat-top light-sheets are very similar, and they both fit closely to the experimental Gaussian data. Simulated flat-top is also performed by the publicly available code from *Remacha et al* (see https://github.com/remachae/beamsimulator) [14].

## Appendix H. Semi-simulated point spread function and optical transfer function

To estimate the point-spread function (PSF) and the optical transfer function (OTF) we multiplied a simulated widefield PSF with the experimentally measured light-sheets. We decided to do it this way because experimentally measured 3D PSFs from beads are often compromised by noise and suffer from background accumulation. In contrast, a simulated PSF is free of any noise. The light-sheet measurements themselves are of high SNR, so we assume that the resulting “semi-simulated” light-sheet PSF has a very high SNR level and thus can faithfully demonstrate the limits of the OTF of each light-sheet system. I.e. in a real-world application, the OTF roll off to the noise level would very likely occur earlier than in our semi-simulated OTFs.

The widefield PSF is generated by the “PSF Generator” plugin in ImageJ [22] using the Born & Wolf 3D Optical Model, and assuming the following parameters: refractive index= 1.33, wavelength= 520 nm, NA= 1.1. Pixel sizes and Z-step were set to 81.25 nm to match our experimental results. The PSFs for the Gaussian light-sheet and the square lattice light-sheet are the product of the widefield PSF with an NA0.31 Gaussian light-sheet and a square lattice with an NA 0.70, na 0.56 annulus, respectively. This particular square lattice was chosen because it is the thinnest square lattice (~0.8 μm) we have created experimentally, and we selected a corresponding Gaussian light-sheet. The optical transfer function (OTF) is calculated by computing the 3D Fourier Transform of the PSF with MATLAB. As can be seen in Figure S6, the OTF for the Gaussian and square lattice light-sheet have a similar sized support, whereas the decay of the Gaussian light-sheet OTF in the kz direction appears more linear. The Lattice light-sheet OTF has two weak sidelobes, corresponding to the interference of the central diffraction orders of the lattice pattern. However, their peak strength is below 3.7% in this ideal, virtually noise free case. Furthermore, there are large gaps between the main body of the OTF and the two sidelobes. As such, we believe that these portions of the OTF unlikely contribute with useful information in a real imaging experiment.

**Fig. S7.**
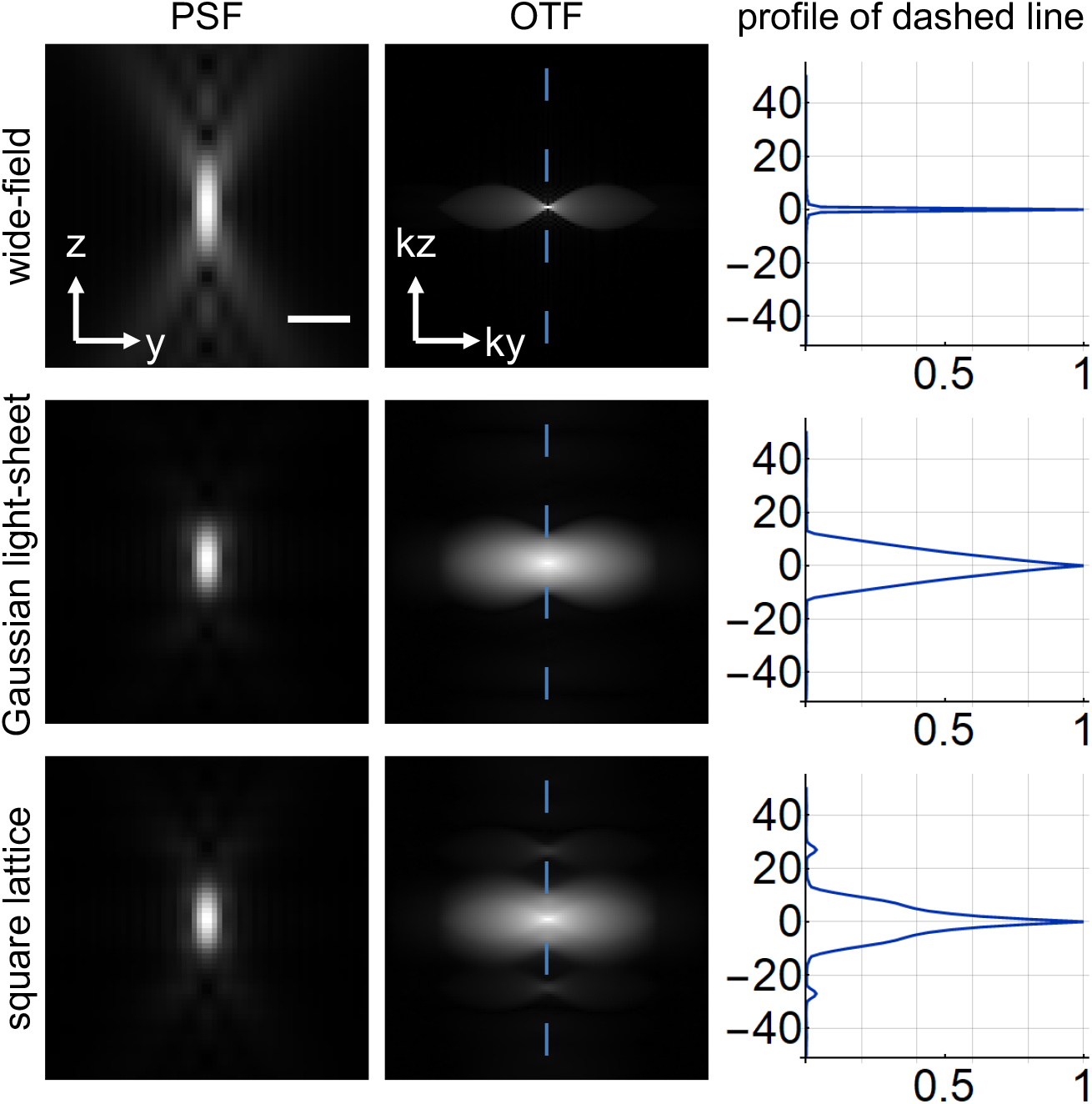
PSFs and OTFs of wide-field, Gaussian light-sheet microscopy, and square lattice light-sheet microscopy. A Gamma correction of 0.5 was applied to all PSF and OTF images to increase the contrast of weak features. Profiles along the dashed lines across the center of OTFs (without the Gamma correction) show clearly the support of the OTFs. Scale bar in the PSFs is 1 μm.

